# Vaginal Microbiome and Inflammation Among Chinese Women

**DOI:** 10.1101/2024.11.30.626174

**Authors:** Zhang Yichan, Schuppe-Koistinen Ina, Wang Huarui, Pelechano Garcia Vicente, Tian Xiangyang, Chen Wei-Hua, Gao Guolan, Du Juan

## Abstract

Human papillomavirus (HPV) infection, cervical intraepithelial neoplasia (CIN), pelvic organ prolapse (POP), cervical polyps (CP), abnormal uterine bleeding (AUB), and vaginitis are common conditions encountered in gynecological practice. In this study, we collected vaginal secretions from patients diagnosed with these diseases (n = 310) and from healthy donors (n = 112) to analyze their vaginal microbiome using 16S rRNA sequencing and cytokine expression using a cytometric bead array (CBA). Our findings revealed that the vaginal microbiome in POP patients exhibited higher complexity compared to other groups. Regarding cytokine expression, levels of IL-1α, IL-1β, IL-6, IL-8, MCP-1, and MIG were significantly elevated in HPV and CIN patients. Analysis of microbial associations showed that *Lactobacillus crispatus* and *Lactobacillus iners* were significantly negatively correlated with IL-1α and IL-1β expression. In contrast, non-*Lactobacillus* bacteria, including *Bifidobacterium breve, Prevotella bivia, Gardnerella vaginalis, Sneathia amnii, Sneathia sanguinegens, Prevotella amnii, Escherichia coli,* and *Chlamydia trachomatis*, were positively correlated with IL-1α, IL-1β, IL-6, IL-8, MCP-1, and MIG levels. Furthermore, *Lactobacillus iners* exhibited a significant negative effect in the HPV and CIN patient groups.

## Introduction

Cervical cancer (CC) is a malignant tumor originating in the cervix [1, 2]. Cervical intraepithelial neoplasia (CIN) is a precancerous lesion strongly associated with the development of invasive cervical cancer and encompasses cervical dysplasia as well as carcinoma in situ. Based on the degree of dysplasia in the cervical squamous epithelium, CIN is classified into three grades: CIN 1, CIN 2, and CIN 3 [3, 4]. Human papillomavirus (HPV), a double-stranded DNA virus, has been identified as a primary causative agent of cervical cancer [5, 6]. HPV types are categorized based on their oncogenic potential into 13 high-risk (HR) types and low-risk (LR) types [7, 8]. High-risk HPV types are closely linked to cervical intraepithelial neoplasia and cervical cancer, while low-risk HPV types are typically associated with mild squamous epithelial injury, urogenital warts, and recurrent respiratory papillomatosis [9, 10, 11, 12].

Pelvic organ prolapse (POP) occurs when the fibromuscular support structures of the pelvic organs weaken or fail, leading to abnormal positioning of the bladder, uterus, rectum, and/or small intestine inside or outside the vagina [13, 14, 15]. POP is a common condition in women, with a prevalence ranging from 30% to 50% among adults, and its incidence increases with age and parity [16, 17, 18]. Cervical polyps (CP) are benign lesions found in 2%–5% of adult women. These growths are thought to result from focal hyperplasia of the glandular epithelium in the endocervix [19, 20, 21]. Abnormal uterine bleeding (AUB) is characterized by changes in menstruation involving increased volume, duration, or frequency of bleeding [22, 23, 24]. This condition is prevalent, affecting 10%–30% of women of reproductive age [25, 26]. Vaginitis accounts for 90% of cases caused by bacterial vaginosis (BV), Candida vaginitis (VVC), or Trichomonas vaginalis (TV) [27, 28, 29, 30, 31, 32]. These infections impact millions of women annually and are associated with adverse outcomes such as preterm labor and delivery, pelvic inflammatory disease (PID), and postabortal endometritis. Other pathogens involved include Neisseria gonorrhoeae, Chlamydia trachomatis, HPV, HSV-2, and HIV-1 [33, 34, 35]. Atrophic vaginitis (AV), a condition affecting 25%–50% of postmenopausal women, is another common affliction [36, 37, 38]. In addition to these diseases, other frequently encountered gynecologic conditions include uterine fibroids, endometrial polyps, ovarian cysts, induced abortion, urinary tract infections (UTIs), cervical lesion treatments, and abnormalities detected via thin-prep cytologic test (TCT) [39, 40, 41, 42, 43, 44].

The vaginal microbiota (VMB) refers to the community of microorganisms present in the vaginal environment of women of reproductive age [45, 46, 47]. The VMB is typically dominated by *Lactobacillus* species, including *Lactobacillus crispatus*, *Lactobacillus iners*, *Lactobacillus gasseri*, and others [48, 49, 50, 51]. These bacteria metabolize glycogen to produce lactic acid, which maintains the vaginal environment at an acidic pH [52, 53, 54]. Research has demonstrated that the vaginal microbiota plays a critical role in the female vaginal microenvironment by supporting vaginal health and serving as the first line of defense against sexually transmitted infections (STIs) [55, 56, 57, 66]. In a healthy state, the vaginal microbiota forms a protective bacterial biofilm on the cervical and vaginal mucosal surfaces [58, 59, 60]. This biofilm secretes lactic acid, bacteriocins, and biological surfactants, which collectively prevent pathogen adhesion, promote autophagy, and facilitate the clearance of harmful bacteria [61, 62, 63].

Abnormal changes in the vaginal microenvironment, often characterized by a predominance of *non–Lactobacillus* species, result in the secretion of short-chain fatty acids (SCFAs) that compromise the protective mechanisms of the vaginal microbiota (VMB) [64, 65, 67]. Such changes disrupt the balance of the VMB by inhibiting the growth of *Lactobacillus* species and increasing the diversity of other microorganisms, particularly strict anaerobic bacteria that produce enzymes and metabolites detrimental to vaginal health [68, 69, 70, 71]. This disruption damages the cervical epithelial barrier and facilitates the development of various diseases [72, 73, 74, 75]. A healthy vaginal microbiome acts as a protective shield against numerous urogenital conditions, including yeast infections, sexually transmitted infections (STIs), urinary tract infections (UTIs), and human immunodeficiency virus (HIV) infection [76, 77, 78, 79]. Conversely, an abnormal microbiome, as observed in bacterial vaginosis (BV), is linked to a heightened risk of upper reproductive tract infections, miscarriage, recurrent miscarriage, preterm birth, HPV infection, and other STIs, including HIV [80, 81, 82, 83].

The immune system plays a crucial role in maintaining the health of the female reproductive tract [84, 85, 86]. The cervix and vagina of adult females harbor a diverse population of immune cells, including T lymphocytes, B lymphocytes, natural killer (NK) cells, dendritic cells (DCs), and mononuclear macrophages, among others [87, 88, 89]. Among these, T lymphocytes are the most abundant, accounting for approximately 50% of all immune cells in the cervix and vagina, and their presence is positively correlated with microbial load [90, 91]. When the state of the cervix changes, the local immune system in the cervix and vagina responds accordingly; however, the specific mechanisms of these changes remain unclear. To investigate this, we initially designed a study to measure 13 immune-related cytokines: GM-CSF, IFN-γ, IL-1α, IL-1β, IL-4, IL-6, IL-8, IL-10, IL-17A, MCP-1, MIG, RANTES, and TNF [92, 93, 94, 95]. In a preliminary experiment involving 100 randomly selected clinical samples, we found that only IL-1α, IL-1β, IL-6, IL-8, MCP-1, and MIG could be reliably quantified. Based on these preliminary results, we proceeded to measure the concentrations of these six cytokines in the remaining samples and analyzed their relationships with vaginal microbiota composition and expression.

In our research, we collected vaginal secretions from clinical patients to analyze their vaginal microbiome and associated cytokine expression. Our goal was to evaluate the expression patterns of these two factors across different gynecological diseases and explore their potential relationships. Since the sequencing results for HPV and CIN samples have already been published, this article focuses on the vaginal microbiome data from five other diseases [96]. Gynecological conditions such as pelvic organ prolapse (POP), cervical polyps (CP), abnormal uterine bleeding (AUB), and various forms of vaginitis are very common in gynecology clinics in China. However, limited research exists on the relationship between the vaginal microbiome (VMB) and these conditions. For diseases like POP, CP, and AUB, the changes in vaginal microbiota associated with these diagnoses remain largely unknown.

Additionally, as part of the vaginal microenvironment, the immune system plays a critical role in cervical and vaginal health. However, studies investigating immune responses in patients with common gynecologic diseases are scarce.

Our study aimed to address this gap by focusing on the relationship between common gynecologic diseases in China, the vaginal microbiome, and the immune system. Notably, our findings revealed that POP exhibits a distinct vaginal bacterial profile compared to other diseases. Furthermore, significant associations were identified between the vaginal microbiome and the cytokines IL-1α and IL-1β.

## Methods

### Study Population

The study was approved by the Aviation General Hospital Ethics Committee, with the certificate number 2021-KY-01-02. The inclusion criteria for participants were as follows: (a) aged between 20 and 70 years; (b) willingness to cooperate during the collection of cervical and vaginal samples; (c) voluntary participation with signed written informed consent; and (d) no plans to move out of Beijing within two years and willingness to cooperate with follow-up assessments. Exclusion criteria included: (a) pregnancy; (b) prior vaccination against HPV; (c) engagement in sexual activity, vaginal irrigation, or drug application within 48 hours before sampling; (d) diagnosis of tuberculosis, hepatitis B virus (HBV), hepatitis C virus (HCV), sexually transmitted infections (e.g., HIV, *Treponema pallidum*), or other infectious diseases; and (e) immune system disorders or chronic diseases such as diabetes mellitus, hypertension, or coronary heart disease.

### Sample Collection and Storage

Clinical samples were collected using two cervical brushes (Huaxia Ruitai Plastic Industry Company, Jiangsu, China). One brush was rinsed in a vial containing normal saline solution for 16S rRNA testing, while the second brush was rinsed in a vial containing PBS solution for cytokine analysis. After sample collection, they were immediately transferred to a −80°C freezer for storage. The samples could be preserved at this temperature for over a month without compromising the integrity of the collected material.

### DNA Extraction

Metagenomic DNA was extracted using the Fast DNA SPIN extraction kits (QIAamp PowerFecal Pro DNA Kit, Qiagen, China). The extraction process involved bacterial DNA being isolated from vaginal secretions via filtration columns followed by rapid centrifugation. Any interference from human tissue DNA was eliminated, and the final extracted DNA sample (100 μL) was obtained. The extracted DNA could also be stored at −80°C for long-term preservation. For sequencing analysis, the DNA samples were sent to Nuohe Zhiyuan Technology Company (Beijing, China).

### 16S rRNA Gene Sequencing

Following DNA extraction, the samples were diluted to 1 ng/μL with sterile water. Specific primers with barcodes were used for PCR amplification. The Phusion High-Fidelity PCR Master Mix with GC Buffer (New England Biolabs, France) was employed for the amplification process. Library preparation was carried out using the TruSeq DNA PCR-free Sample Preparation Kit (Illumina Inc., San Diego, USA), followed by library quantification using Qubit and q-PCR. The prepared library was then sequenced using the NovaSeq high-throughput sequencing platform (PE250, Illumina Inc., San Diego, USA).

### Sequencing Data Analysis

After amputation of barcode and primer sequences, reads from each sample were merged using FLASH (V1.2.7) (http://ccb.jhu.edu/software/FLASH/) to obtain the Raw Tags. These raw tags were then subjected to stringent filtering to generate high-quality Clean Tags. The quality control process followed the Reference Qiime Tags approach, and the final Effective Tags were obtained using the Uparse algorithm for all samples (http://drive5.com/uparse/). Sequences were clustered into operational taxonomic units (OTUs) with 97% identity by default. The OTU sequences were annotated by species using the Mothur method and the SSUrRNA database of SILVA138. For multiple sequence alignment, the MUSCLE software (https://www.drive5.com/muscle/) was used. The Qiime software (Version 1.9.1) was employed to calculate OTUs, Chao1, Shannon, Simpson, and beta-diversity analyses. Additionally, R software (Version 2.15.3) was used to generate dilution curves, rank abundance curves, species accumulation curves, and Principal Coordinates Analysis (PCoA) plots.

### Cytometric Bead Array (CBA)

The standard freeze-dried microspheres were poured into the same test tube, and an Assay Diluent was used to dilute the standard material. After standing for 15 minutes, the material underwent multiple dilutions according to a 2n factor. Capture Beads were then mixed and added to the flow tubes. Equal amounts of both standard and sample were added to each tube. The tubes were mixed well and incubated at room temperature away from light for 1 hour. After incubation, a buffer was added to clean excess antibodies, and the antibody solution was suspended for testing using a cytometric bead array machine.

### Co-culture and Supernatant Collection

Hela cells were cultured in Dulbecco’s Modified Eagle’s Medium (DMEM) with high glucose (Sigma, Cat. No: D6429-500ml) and supplemented with 10% fetal bovine serum (FBS-HI) (Sigma, Cat. No: F9665-500ml). The cells were incubated at 37°C with 5% CO2. Bacteria used in co-culture experiments were purchased from CCUG (Culture Collection University of Gothenburg) and cultured in NYCIII medium containing 10% FBS. The strains used for co-culture with Hela cells included: Lactobacillus crispatus (CCUG 38700), Gardnerella vaginalis (CCUG 44012), Prevotella bivia (CCUG 34046), Lactobacillus iners (CCUG 44108), Bifidobacterium longum (CCUG 59493), Aerococcus christensenii (CCUG 28831T), Fannyhessea vaginae “V” (CCUG 44116), Sneathia sanguinegens (CCUG 42621), Sneathia amnii (CCUG 52977T), Lactobacillus gasseri (CCUG 44082), Bifidobacterium breve (CCUG 27171A), Escherichia coli (CCUG 44113).

For the co-culture experiment, 2 × 10⁵ Hela cells were plated into a 12-well plate and incubated at 37°C, 5% CO2. After overnight incubation, cells were washed with Dulbecco’s Phosphate Buffered Saline (DPBS) (Gibco, Cat. No: 14200-067, 500ml), and 500 μL of DMEM without FBS was added to keep the cells moist. Bacterial cultures were diluted in DMEM, and the bacterial OD (optical density) was measured and adjusted to the appropriate concentration. The bacteria (500 μL) were added to the Hela cells at a MOI (multiplicity of infection) of 10. After 6 and 24 hours of co-culturing, supernatants were collected and stored in tubes at −80°C for later analysis.

### ELISA Test

The ELISA (Enzyme-Linked Immunosorbent Assay) was used to measure the cytokine concentrations in the supernatants. Reagents and antibodies for ELISA were purchased from Thermo Fisher. The following capture antibodies were used: IL-6 (Monoclonal Antibody, PeproTech®, Fisher Scientific, Cat. No: 500-M06-500UG). IL-8 (Monoclonal Antibody, PeproTech®, Fisher Scientific, Cat. No: 500-M08-500UG). IL-1 beta (Monoclonal Antibody, PeproTech®, Fisher Scientific, Cat. No: 500-M01B-500UG). IL-1 alpha (Monoclonal Antibody (ILA9-H18), Fisher Scientific, Cat. No: M421A). The corresponding detection antibodies were: IL-6 (Polyclonal Antibody, Biotin, PeproTech®, Fisher Scientific, Cat. No: 500-P26GBT-50UG). IL-8 (Polyclonal Antibody, Biotin, PeproTech®, Fisher Scientific, Cat. No: 500-P28BT-50UG). IL-1 beta (Polyclonal Antibody, Biotin, PeproTech®, Fisher Scientific, Cat. No: 500-P21BGBT-50UG). IL-1 alpha (Monoclonal Antibody (ILA8-H12), Biotin, Fisher Scientific, Cat. No: M420AB). The ELISA plates were measured using the SpectraMax MiniMax™ 300 Imaging Cytometer with a 450nm laser, and the results were analyzed using the supporting software.

### Statistical Analysis

The Kruskal-Wallis test was employed to compare microbial alpha and beta diversities between two or more groups. Benjamini-Hochberg correction was applied to adjust for multiple testing and reduce the false discovery rate. The relationship between HPV infection, vaginal microbiota, and age was assessed using the χ² (Chi-square) test. All statistical analyses were conducted using SPSS version 27 (IBM, New York, NY) and R software version 3.6.3. A P value of less than 0.05 was considered statistically significant. Graphical images were generated using GraphPad Prism 9.0. This methodological approach ensures rigorous statistical analysis and accurate interpretation of results.

## Results

### Participant Characteristics in the Study Cohort

The participants in this study were Chinese women who visited the Aviation General Hospital of China Medical University from November 2020 to May 2021. After excluding unqualified samples, 422 valid samples were included in the study. These participants were divided into 8 groups: a healthy control group (n=112), an HPV-positive group (n=156), a CIN-diagnosed group (n=97), a pelvic organ prolapse (POP) group (n=9), a cervical polyp (CP) group (n=6), an abnormal uterine bleeding (AUB) group (n=8), a vaginitis group (n=20), and an “other” group (n=14) (Supplementary Overview Figure).

### Significant Differences in Sequencing Results Among the 8 Disease Groups

The POP group exhibited the highest species diversity and was distinctively different from the CP, AUB, vaginitis, and other groups. Compared with the other groups, POP had a lower percentage of *Lactobacillus crispatus* and *Lactobacillus iners*, but a higher percentage of other species (Figures 1A and B). In alpha diversity analysis, the POP group showed significantly higher Simpson diversity index values compared to the healthy control group, indicating a much larger diversity of species (Figure 1C). Additionally, beta-diversity analysis using PCoA revealed that the POP group was farthest from the other groups, indicating a significant difference in microbial composition (Figure 1D and E).

**Figure 1:**
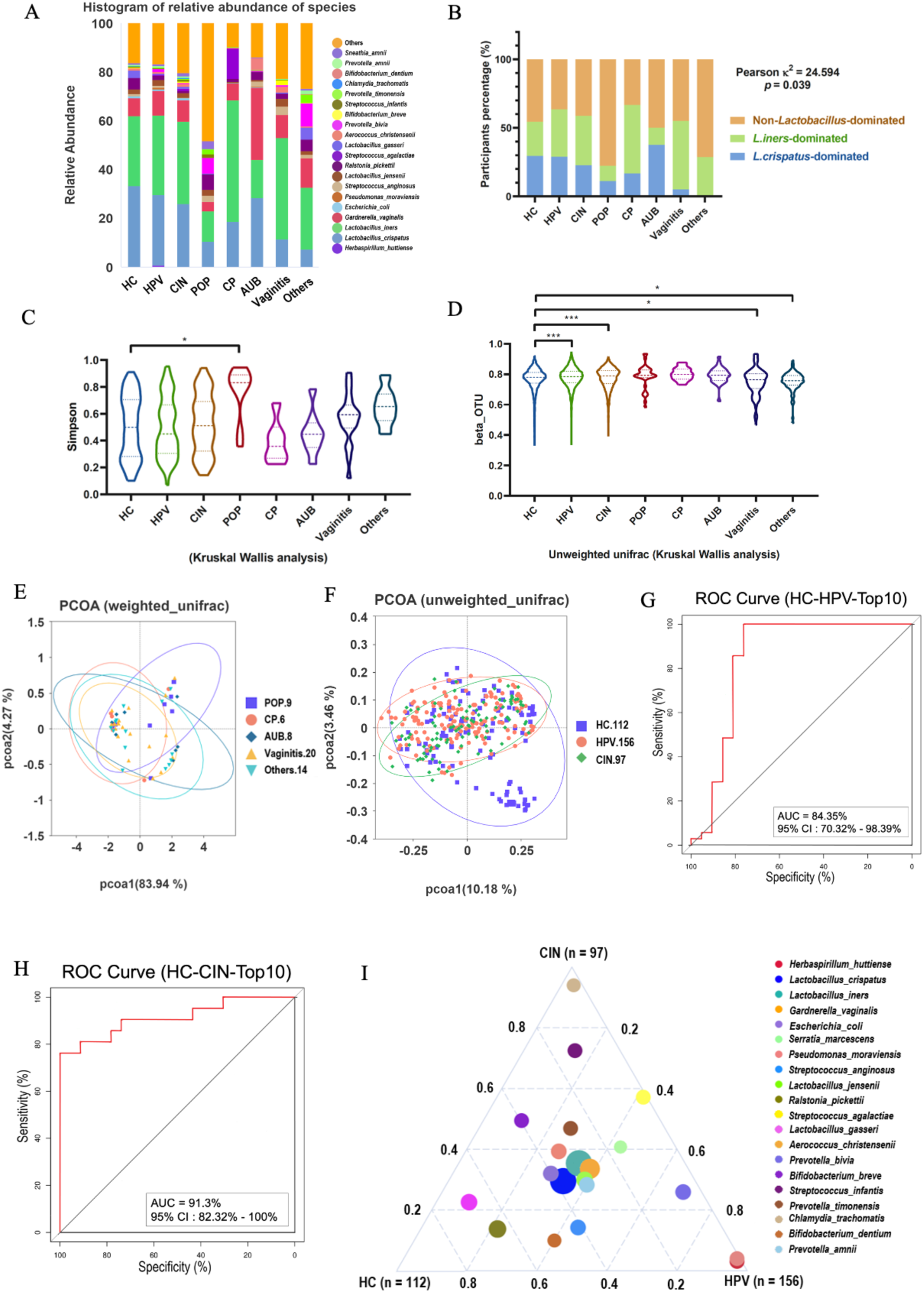
Sequencing results of clinical samples. **A)** Histogram of relative abundance in 8 groups which included HPV and CIN groups. **B)** Bar chart of participants percentage in 8 groups, which was divided by *L.crispatus* – dominated, *L. iners* – dominated, and Non – *Lactobacillus* – dominated. **C)** Alpha analysis of healthy control, POP, CP, AUB, Vaginitis, and others group. **D)** Beta analysis of OUT in 8 groups, which used unweighted unifrac distance algorithm. **E)** PCoA results of POP, CP, AUB, Vaginitis and others groups under weighted unifrac distance algorithm. **F)** PCoA results of HC, HPV, and CIN groups under unweighted unifrac distance algorithm. **G)** ROC curve of HC and HPV groups in top 10. **H)** ROC curve of HC and CIN groups in top 10. **I)** Terneryplot analysis of HC, HPV, and CIN groups. (Kruskal Wallis analysis, * : p < 0.05; ** : p < 0.01; *** : p < 0.001)

In comparison with the healthy control group, both the HPV and CIN groups exhibited a lower percentage of *Lactobacillus crispatus* and *Lactobacillus iners*, and a higher percentage of *Gardnerella vaginalis* and other non-*Lactobacillus* bacteria, as shown in the bar chart of species percentages (Figures 1A and B). Since CIN represents the advanced stage of HPV, the CIN group displayed an even lower percentage of *Lactobacillus crispatus* and *Lactobacillus iners* than the HPV group (Figures 1A and B). In beta-diversity analysis, both HPV and CIN groups also showed significantly higher diversity than the healthy control group (Figure 1D). However, there was no clear distinction among the HC, HPV, and CIN groups in unweighted UniFrac PCoA analysis (Figure 1F). For alpha diversity, both the HPV and CIN groups exhibited higher Shannon and Chao1 indices compared to the control group. Since these results have already been published in our previous article, we did not include them in this report.

For further analysis, we conducted random forest analysis on the HC, HPV, and CIN groups and identified the top 10 species based on Mean Decrease Accuracy (Supplementary Figures 1E and F). Based on the results of the random forest analysis, we constructed ROC (Receiver Operating Characteristic) curves using cross-validation. The AUC for the HC vs. HPV comparison was 84.35% (Figure 1G), and for the HC vs. CIN comparison, the AUC was 91.3% (Figure 1H). These AUC values indicate that the classification model performed well. To further investigate differences in dominant species between the HC, HPV, and CIN groups, we analyzed the top 20 species by average abundance and visualized the data using a ternary plot (Figure 1I). The dominant species in the HC group were *Lactobacillus gasseri* and *Ralstonia pickettii*, while in the HPV group, the dominant species were *Bifidobacterium dentium* and *Escherichia coli*, and in the CIN group, the dominant species were *Bifidobacterium breve* and *Chlamydia trachomatis*. Other species, such as *Lactobacillus crispatus*, *Lactobacillus iners*, and *Gardnerella vaginalis*, were found in the intermediate positions, showing no clear dominance (Figure 1I).

To classify the groups further, we divided the HC, HPV, and CIN groups into sub-groups dominated by *Lactobacillus crispatus*, *Lactobacillus iners*, or Non–*Lactobacillus* species. PCoA analysis of the 9 sub-groups revealed that the three *L. crispatus*-dominated sub-groups clustered together, as did the *L. iners*-dominated and non–Lactobacillus–dominated sub-groups (Supplementary Figure 1D).

### Cytokine Expression Differences in HC, HPV, and CIN Groups

We tested the expression levels of six inflammation-related cytokines in all clinical samples, including IL-1α, IL-1β, IL-6, IL-8, MCP-1, and MIG. Among these, MIG and IL-8 showed significant differences across the eight disease groups (Supplementary Figures 2A and D). Due to the much larger sample sizes in the HC, HPV, and CIN groups compared to the other five groups, which could introduce statistical bias, we focused on these three groups for further analysis. In the HPV group, MIG concentration was significantly higher than in the healthy control group (Figure 2F). In the CIN group, concentrations of IL-1α, IL-1β, IL-8, and MCP-1 were notably elevated compared to the HPV group (Figures 2A, B, D, and E). Moreover, the concentrations of IL-1α and MIG in the CIN group were much higher than those in the control group (Figures 2A and F).

**Figure 2:**
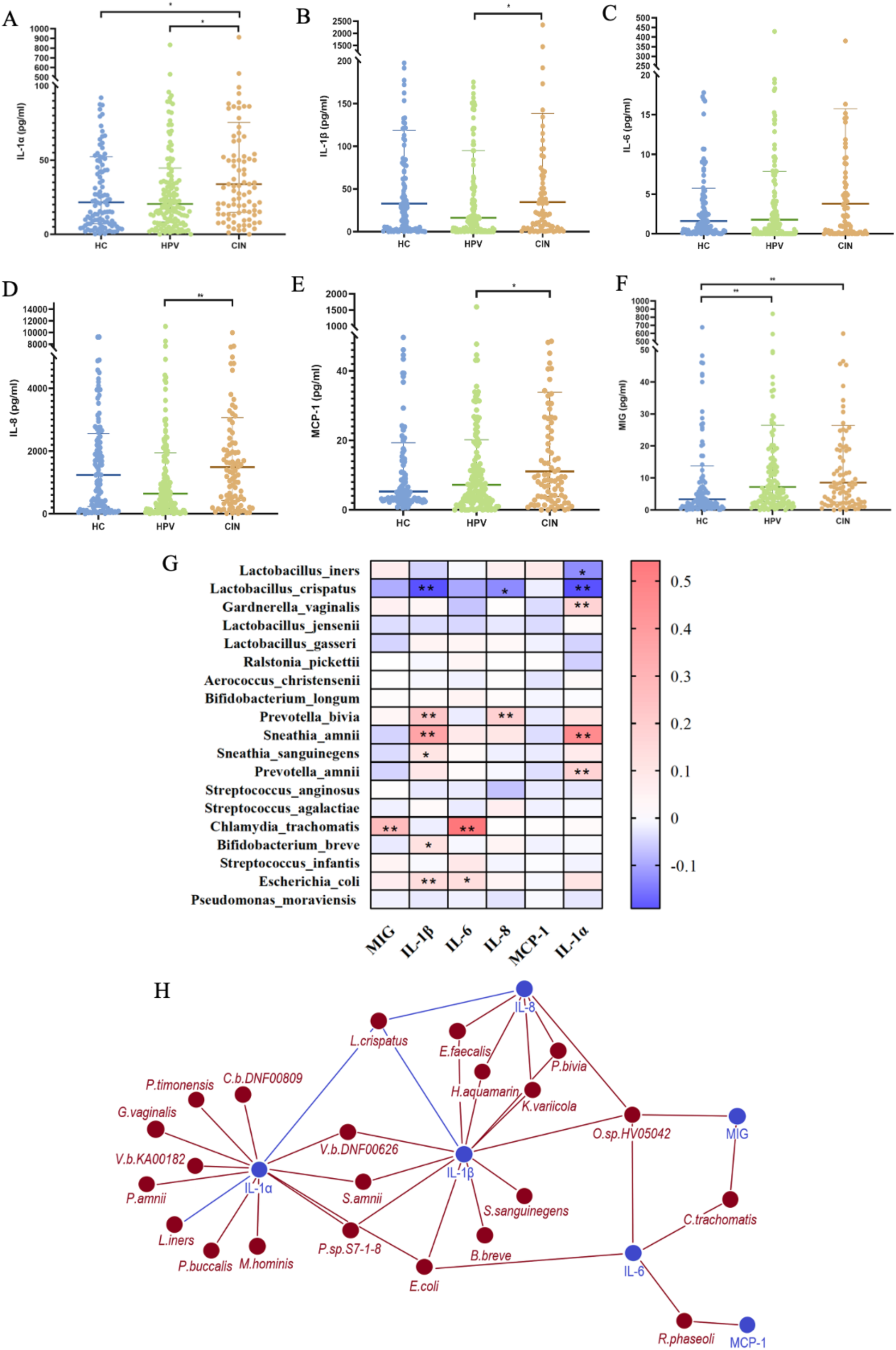
Cytokines and bacteria expression in HC, HPV and CIN groups. **A-F)** Concentrations of IL-1α, IL-1β, IL-6, IL-8, MCP-1 (monocyte chemoattractant protein) and MIG (monokine induced by gamma-interferon) showed in HC, HPV and CIN groups. (Kruskal Wallis analysis, * : p < 0.05; ** : p < 0.01; *** : p < 0.001) **G)** Heatmap of 6 cytokines and top 19 strains. **H)** Network of top 50 bacteria which have statistic significant with related cytokines. (Pearson analysis, * : p < 0.05; ** : p < 0.001)

### Positive and Negative Relationships Between Cytokines and Bacteria

We also investigated the relationships between bacterial species and cytokine expression. A heatmap was created to illustrate these positive and negative correlations between the top 19 bacterial strains and the six cytokines (Figure 2G). While no statistically significant relationship was observed between MCP-1 and the strains, several notable correlations were found. *Chlamydia trachomatis* and MIG have positive relationship. *Bifidobacterium breve, Sneathia amnii, Sneathia sanguinegens, Prevotella bivia* and *Escherichia coli* have positive relationship with IL-1β. While *Lactobacillus crispatus* has negative relationship with IL-1β. *Sneathia amnii, Prevotella amnii,* and *Gardnerella vaginalis* have positive relationship with IL-1α, while *Lactobacillus crispatus* and *Lactobacillus iners* have negative relatoinship with IL-1α. *Escherichia coli* and *Chlamydia trachomatis* have positive relationship with IL-6. *Prevotella bivia* has positive relationship with IL-8, while *Lactobacillus crispatus* has negative relationship with IL-8 (Figure 2G). We also showed their relationship using network analysis, and red lines showed positive relationship, while blue lines showed negative relationship (Figure 2H).

### Cytokine Expression and Its Relationship with Bacterial Species

As previously described, we divided the HC, HPV, and CIN groups into three subgroups based on the dominant bacterial species: *L. crispatus*-dominated, *L. iners*-dominated, and MIX (*Non-Lactobacillus*-dominated). This resulted in a total of nine subgroups. We then compared cytokine expression among these subgroups within the same disease group. In the HC group, significant differences in cytokine levels were only observed for IL-1β. Specifically, the *L. iners*-dominated and MIX sub-groups exhibited significantly higher IL-1β levels than the *L. crispatus*-dominated subgroup (Figure 3A). No other cytokines showed significant differences between the subgroups (Supplementary Figures 3A, B, C, D, and E). The PCA plot indicated a relationship between IL-1β and bacteria such as *L. iners* and *Escherichia coli*, as these strains clustered in the same quadrant and direction as IL-1β (Figure 3B).

**Figure 3:**
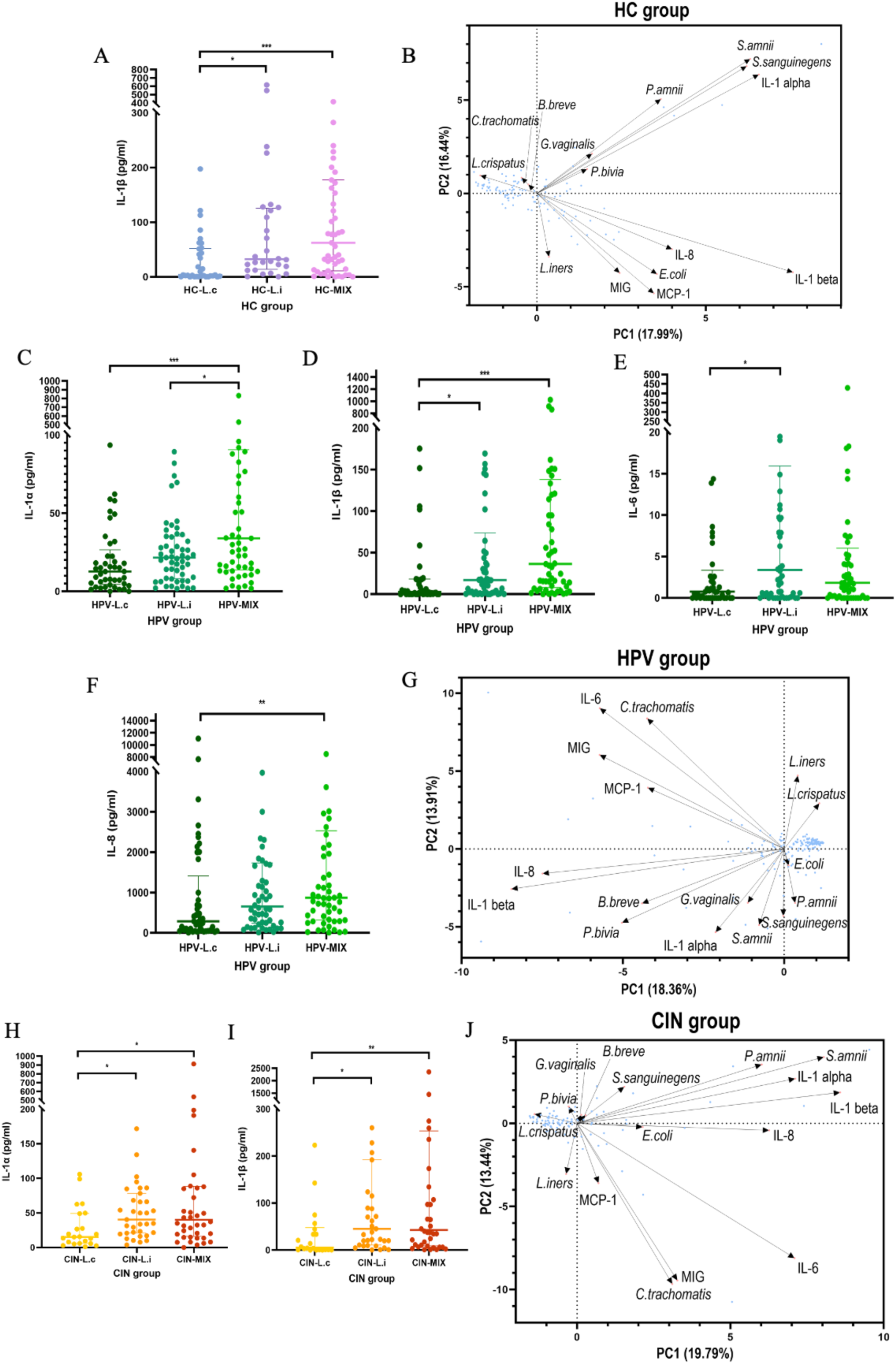
Scatter plot and Principal component analysis (PCA). **A-B)** IL-1β has relationship with bacteria in HC group. **C-G)** IL-1α, IL-1β, IL-6 and IL-8 have relationship with strains in HPV group. **H-J)** IL-1α and IL-1β have relationship with bacteria in CIN group. (Kruskal Wallis analysis, * : p < 0.05; ** : p < 0.01; *** : p < 0.001)

In the HPV group, the MIX subgroup had significantly higher concentrations of IL-1α, IL-1β, and IL-8 compared to the *L. crispatus*-dominated subgroup (Figures 3C, D, and F). In contrast, IL-1β and IL-6 concentrations were significantly higher in the *L. iners*-dominated subgroup compared to the *L. crispatus*-dominated subgroup (Figures 3D and E). Additionally, the concentration of IL-1α in the MIX subgroup was higher than in the *L. iners*-dominated subgroup (Figure 3C). No significant differences were found in the expression of MCP-1 and MIG across these three subgroups (Supplementary Figures 3F and G). In the PCA plot, IL-6 clustered with *Chlamydia trachomatis*, while IL-1α, IL-1β, and IL-8 were associated with species such as *Bifidobacterium breve, Prevotella bivia, Gardnerella vaginalis,* and *Sneathia amnii* (Figure 3G). In the CIN group, the *L. iners*-dominated and MIX sub-groups showed significantly higher concentrations of IL-1α and IL-1β compared to the *L. crispatus*-dominated subgroup (Figures 3H and I). However, no significant changes were observed in IL-6, IL-8, MCP-1, and MIG (Supplementary Figures 3H, I, J, and K). The PCA plot indicated that *Bifidobacterium breve, Gardnerella vaginalis, Sneathia amnii, Sneathia sanguinegens,* and *Prevotella amnii* were associated with IL-1α and IL-1β (Figure 3J).

### Comparison of Cytokine Expression in Subgroups Across Different Diseases

As shown in Supplementary Figure 1D, bacterial profiles for the *L. crispatus*-dominated subgroup in HC, HPV, and CIN groups clustered together, with similar trends observed for the *L. iners*-dominated and MIX groups. We further analyzed the cytokine concentrations within these subgroups across the three disease groups. In the *L. crispatus*-dominated and MIX sub-groups, no significant differences in cytokine concentrations were found across the HC, HPV, and CIN subgroups (Supplementary Figure 4). However, in the *L. iners*-dominated subgroup, the concentrations of IL-1α and IL-8 were significantly higher in the CIN subgroup compared to the HPV subgroup (Supplementary Figures 5A and B). Additionally, MIG expression was higher in the CIN subgroup compared to the HC subgroup (Supplementary Figure 5C).

### Positive and Negative Relationships Between Bacterial Species and Cytokines

We analyzed the heatmap and network of bacteria and cytokines in HC, HPV, and CIN groups and tried to find out their difference. In HC group, *Lactobacillus crispatus* has significantly negative relationship with IL-1α and IL-1β (Figure 4B). IL-1α has significantly positive correlation with *Prevotella amnii, Sneathia amnii,* and *Sneathia sanguinegens*. IL-6 has obviously positive correlation with *Escherichia coli*, *Lactobacillus gasseri*, and *Bifidobacterium longum*. IL-1β has significantly positive relationship with *Escherichia coli* and *Sneathia amnii.* MIG has significantly positive relationship with *Escherichia coli* (Figure 4A, and B).

**Figure 4:**
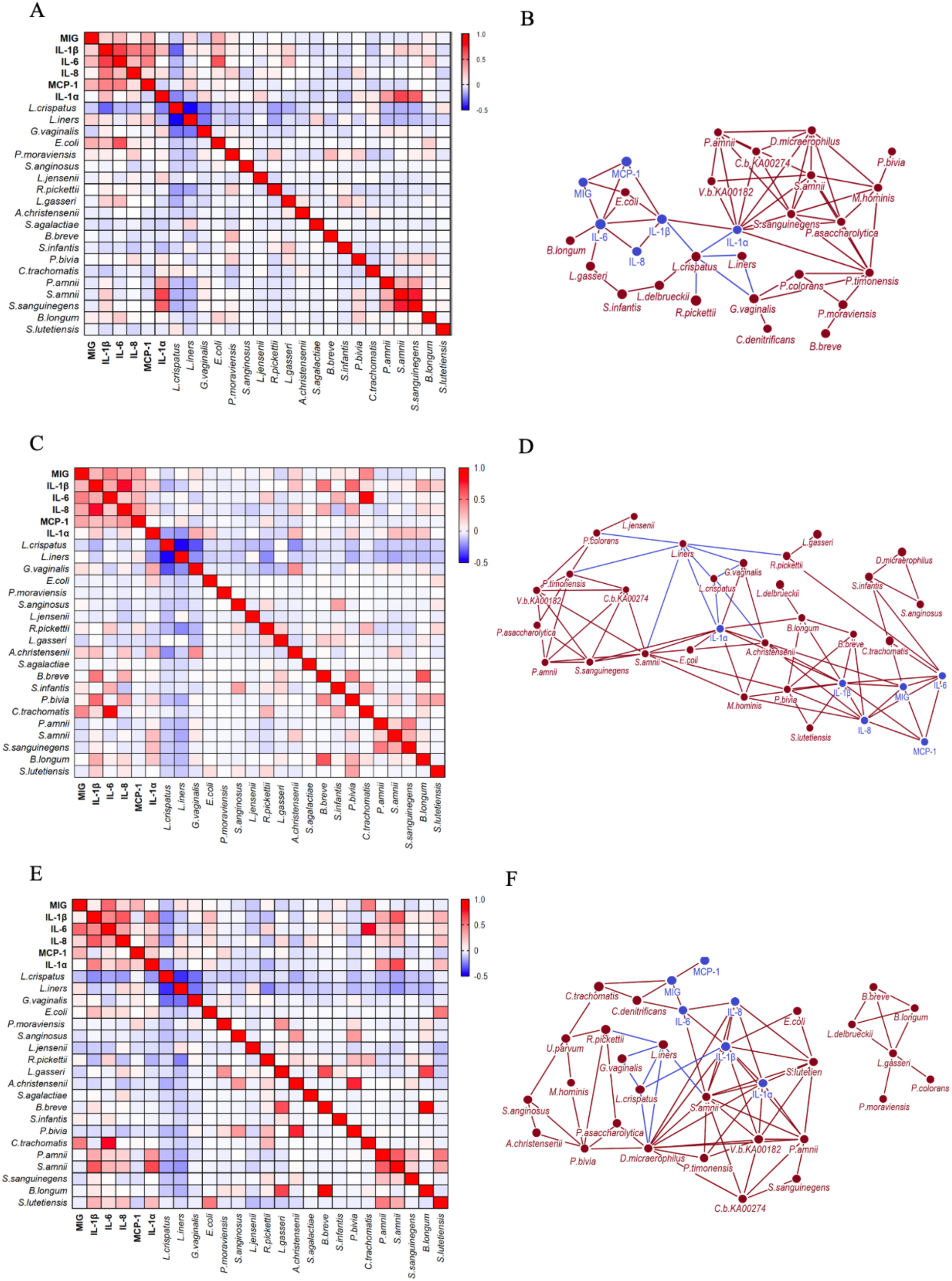
Pearson Heatmap and network in HC, HPV and CIN groups. **A)** Heatmap of 6 cytokines and top 20 strains in HC group. **B)** Network of top 30 bacteria which have statistic significant with related cytokines in HC group. **C)** Heatmap of 6 cytokines and top 20 strains in HPV group. **D)** Network of top 30 bacteria which have statistic significant with related cytokines in HPV group. **E)** Heatmap of 6 cytokines and top 20 strains in CIN group. **F)** Network of top 30 bacteria which have statistic significant with related cytokines in CIN group. (Pearson analysis, * : p < 0.05; ** : p < 0.001)

In HPV group, IL-1α has obviously negative correlation with *Lactobacillus crispatus* and *Lactobacillus iners* (Figure 4D), and has significantly positive correlation with *Gardnerella vaginalis, Escherichia coli, Aerococcus christensenii, Sneathia amni*i, *Sneathia sanguinegens*, and *Bifidobacterium longum.* IL-8 has significantly positive correlation with *Aerococcus christensenii, Bifidobacterium breve,* and *Prevotella bivia.* IL-6 has obviously positive relationship with *Ralstonia pickettii*, *Streptococcus infantis,* and *Chlamydia trachomatis.* IL-1β has obviously positive correlation with *Aerococcus christensenii, Bifidobacterium breve, Prevotella bivia, Bifidobacterium longum,* and *Streptococcus lutetiensis*. MIG has obviously positive correlation with *Aerococcus christensenii* and *Chlamydia trachomatis* (Figure 4C, and D).

In CIN group, only IL-1β has negative relationship with *Lactobacillus crispatus,* and has obviously positive correlation with *Escherichia coli, Prevotella amnii, Sneathia amni*i, and *Streptococcus lutetiensis* (Figure 4F). IL-1α has obviously negative correlation with *Prevotella amnii, Sneathia amni*i, and *Streptococcus lutetiensis.* IL-8 has significantly positive correlation with *Sneathia amnii,* and *Streptococcus lutetiensis.* IL-6 has obviously positive relationship with *Chlamydia trachomatis* and *Sneathia amnii.* MIG has obviously positive correlation with *Chlamydia trachomatis* (Figure 4E, and F).

### Cytokine Expression in Co-cultured Supernatant Compared to Clinical Samples

We co-cultured Hela cells with different bacterial species and measured cytokine levels (IL-1α, IL-1β, IL-6, IL-8) in the supernatant after 6 and 24 hours of co-culturing. The cytokine patterns observed in these experiments were similar to those in the clinical samples. At 6 hours, we observed that *Gardnerella vaginalis*, *Prevotella bivia, Sneathia amnii*, and *Sneathia sanguinegens* significantly increased the levels of IL-1β and IL-1α, while *Lactobacillus gasseri* and *Aerococcus christensenii* showed a decrease in IL-1β. On the other hand, *Lactobacillus crispatus, Sneathia sanguinegens,* and *Fannyhessea vaginae* caused lower IL-8 expression, while *Lactobacillus gasseri* and *Sneathia amnii* showed increased IL-8 levels. For IL-6, *Lactobacillus crispatus, Gardnerella vaginalis*, and *Prevotella bivia* significantly lowered its concentration, while *Lactobacillus iners* also reduced IL-6 at 6 hours (Figure 5A, C, E and G).

**Figure 5:**
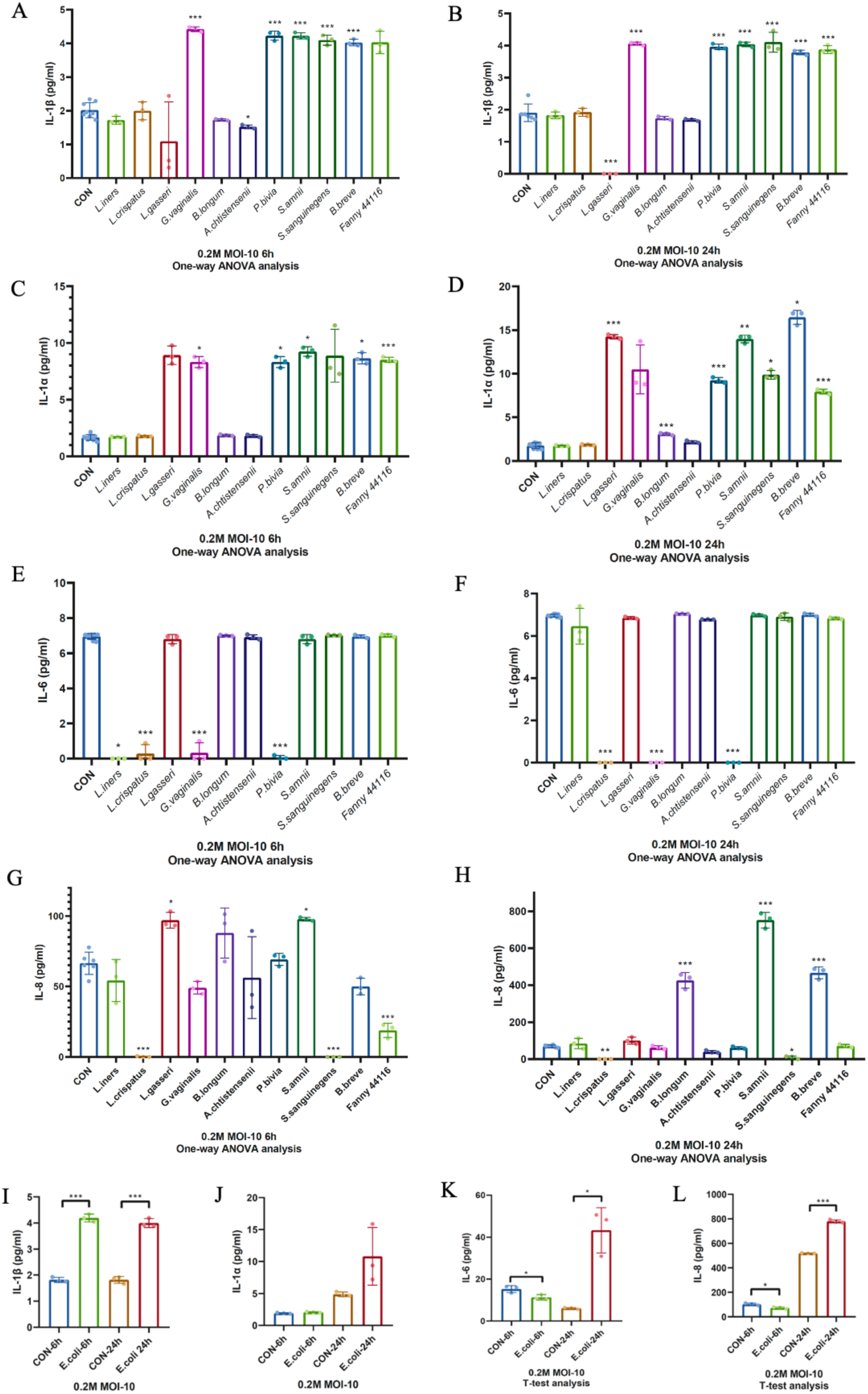
Cytokines concentration tested by ELISA. **A-B)** IL-1β in supernatant co-cultured after 6 and 24 hours. **C-D)** IL-1α in supernatant co-cultured after 6 and 24 hours. **E-F)** IL-6 in supernatant co-cultured after 6 and 24 hours. **G-H)** IL-8 in supernatant co-cultured after 6 and 24 hours. **I-L)** IL-1β, IL-1α, IL-6, and IL-8 in supernatant which collected from *Escherichia_coli* and Hela cells co-culturation after 6 and 24 hours. (One-way ANOVA analysis, * : p < 0.05; ** : p < 0.01; *** : p < 0.001; T-test analysis, * : p < 0.05; ** : p < 0.01; *** : p < 0.001)

At 24 hours, the cytokine responses continued to vary. *Gardnerella vaginalis, Sneathia amnii,* and *Fannyhessea vaginae* induced a significant increase in IL-1β and IL-1α levels, while *Lactobacillus gasseri* and *Bifidobacterium longum* showed lower expression for IL-1α. IL-8 levels were significantly higher for *Bifidobacterium longum, Sneathia amnii*, and *Bifidobacterium breve*, while *Lactobacillus crispatus* and *Sneathia sanguinegens* caused a decrease in IL-8. Additionally, *Lactobacillus crispatus, Gardnerella vaginalis*, and *Prevotella bivia* significantly reduced IL-6, and *Lactobacillus iners* continued to show lower levels at 24 hours (Figure 5B, D, F and H).

For Escherichia coli, which was co-cultured under a microoxygen environment, IL-1β showed a significant increase at both 6 and 24 hours. However, IL-6 and IL-8 exhibited opposing patterns: at 6 hours, their levels were lower than the control, while at 24 hours, they were significantly higher. No significant changes in IL-1α were observed with *Escherichia coli* co-culture (Figure 5I, J, K, and L).

In summary, bacterial species showed varying impacts on cytokine expression, with some bacteria, such as *Gardnerella vaginalis, Sneathia amnii,* and *Fannyhessea vaginae*, increasing IL-1β, IL-1α, and IL-8 levels, while others like *Lactobacillus crispatus* and *Prevotella bivia* decreased IL-6 levels. These patterns observed in the co-culture experiment closely resembled those from the clinical samples, suggesting that the bacteria present in the vaginal microbiome may influence immune responses in gynecological diseases.

## Discussion

### Overview of Clinical Patient Sample Collection and Vaginal Microbiome Analysis

In this study, we collected vaginal secretion samples from patients with various gynecological conditions to analyze their vaginal microbiomes and the expression of related cytokines. The patients were diagnosed with a range of conditions including HPV infection, CIN (Cervical Intraepithelial Neoplasia) 1, 2, and 3, pelvic organ prolapses (POP), cervical polyps (CP), abnormal uterine bleeding (AUB), vaginitis, and other gynecological diseases (Supplementary Overview Figure). Based on these diagnoses, the patients were divided into different groups for analysis. Since our focus was on colposcopy clinic patients, the majority of the samples came from those diagnosed with HPV infection and CIN. We have previously published the alpha-diversity analysis results for these groups, so this article does not repeat that data. Instead, we focus on the analysis of the POP, CP, AUB, vaginitis, and other groups (Figure 1C, Supplementary Figures 1A and B). In terms of beta-OTU (Operational Taxonomic Unit) analysis, the HPV and CIN groups showed significantly higher diversity compared to the healthy control group, highlighting a clear difference between these three groups (Figure 1D, Supplementary Figure 1C).

### Unique Findings in the POP Group

The POP (Pelvic Organ Prolapse) group displayed distinct characteristics compared to the other patient groups. In terms of alpha-diversity (a measure of microbial richness and evenness), the Shannon and Simpson indices for POP patients were significantly higher than those for the healthy control group, indicating that the vaginal microbiome of POP patients was more diverse (Figure 1C, Supplementary Figure 1A, and B). In addition to the diversity measures, the microbial composition of the POP group was notably different. The proportion of *Lactobacillus crispatus* and *Lactobacillus iners*, which are typically abundant in a healthy vaginal microbiome, was significantly reduced in the POP group. Conversely, the percentage of *non-Lactobacillus* bacteria was higher in these patients (Figure 1A and B). The Principal Coordinate Analysis (PCoA) plot further emphasized this trend, showing that the POP group’s microbial profile clustered separately from the other groups, including those with HPV, CIN, CP, AUB, and vaginitis (Figure 1E).

It is worth noting that electrical stimulation therapy is a common treatment for POP patients. During this treatment, an electrode tip is inserted into the vagina, which could potentially influence the stability of the vaginal microbiome, possibly by disrupting the bacterial biofilm. This intervention might explain the observed higher alpha-diversity, reduced percentage of *Lactobacillus spp*., and increased presence of *non-Lactobacillus* bacteria in the vaginal microbiomes of POP patients. These findings suggest that treatment-related factors, such as electrical stimulation, may impact the vaginal microbial composition and contribute to the altered microbiome observed in POP patients.

### Cytokine Expression and Its Correlation with Vaginal Microbiome in HC, HPV, and CIN Groups

We observed significant differences in the expression of several inflammation-related cytokines—IL-1α, IL-1β, IL-8, MCP-1, and MIG—among the healthy control (HC), HPV-infected, and CIN-diagnosed groups. These cytokines are related to immune response and inflammation, and we found that their concentrations in the CIN group were significantly higher than in the HPV or healthy control groups, suggesting a heightened inflammatory state in the vaginal microenvironment of CIN patients (Figure 2A-F). However, we did not observe significant changes in cytokine expression in the other clinical groups such as POP, CP, AUB, Vaginitis, and Others (Supplementary Figure 2A-F), so our analysis continued to focus on HPV and CIN groups. The ternary plot analysis of the vaginal microbiome in the HC, HPV, and CIN groups revealed that each group had a distinct dominant bacterial species (Figure 1I). Based on this, we hypothesized that certain bacterial species may correlate with the expression of inflammation-related cytokines. To explore this, we performed a random forest analysis comparing the HC and HPV groups, identifying the top 10 bacterial species in each group (Supplementary Figure 1E and F). Additionally, we categorized the patients into three sub-groups based on the dominance of specific bacteria: *Lactobacillus crispatus*-dominated, *Lactobacillus iners*-dominated, and *Non-Lactobacillus-*dominated (MIX) sub-groups. These sub-groups clustered distinctly in Principal Coordinates Analysis (PCoA) (Supplementary Figure 1D).

To understand the relationship between vaginal bacteria and cytokines, we analyzed the heatmap and network of cytokine and bacterial species interactions across all patient samples (Figure 2G, and H). The results revealed that IL-1α and IL-1β were correlated with a broader range of bacterial species than the other cytokines, indicating that they play a central role in the vaginal microenvironment (Figure 2G). Both the heatmap and network analysis showed that *Lactobacillus crispatus* and *Lactobacillus iners* were negatively correlated with cytokines, suggesting that these *Lactobacillus* species may have an inhibitory effect on inflammation within the vaginal environment. In contrast, several *non-Lactobacillus* species, including *Bifidobacterium breve, Prevotella bivia, Gardnerella vaginalis, Sneathia amnii, Sneathia sanguinegens, Prevotella amnii, Escherichia coli,* and *Chlamydia trachomatis*, exhibited a positive correlation with cytokine levels, indicating that these bacteria may contribute to the amplification of inflammation in the vaginal microenvironment (Figure 2G, and H).

To further investigate the relationship between bacterial composition and cytokine expression, we compared the cytokine levels across the three sub-groups (*L. crispatus-* dominated, *L. iners*-dominated, and *Non-Lactobacillus*-dominated (MIX)) in each disease group. We found that cytokine concentrations in the *L. iners*-dominated and MIX sub-groups were significantly higher than in the *L. crispatus*-dominated sub-group across all three groups (HC, HPV, and CIN) (Figure 3A, C, D, E, F, H, and I). This result suggests that *L. iners*-dominated and MIX sub-groups are associated with increased inflammation, regardless of whether the samples were from the healthy control, HPV, or CIN groups. The PCA plot provided additional evidence of a relationship between specific cytokines and bacterial species, as certain cytokines and bacterial species clustered in the same quadrant, indicating a correlation between cytokine expression and the vaginal microbiome (Figure 3B, G, and J). This supports the idea that cytokines and the vaginal microbiome are interrelated and that *non-Lactobacillus* bacteria may contribute to inflammation in the vaginal environment, particularly in the *L. iners*-dominated and MIX sub-groups. Together, these findings underscore the importance of the vaginal microbiome in regulating inflammation and highlight the potential role of specific bacterial species in influencing cytokine expression and the inflammatory state of the vaginal microenvironment.

### *Lactobacillus iners* and Its Inhibitory Role in HPV and CIN Groups

We further analyzed the correlation between bacterial species and cytokine expression in the HC, HPV, and CIN groups using heatmap and network analyses (Figure 4). *Lactobacillus crispatus* consistently showed a negative correlation with various bacteria, suggesting that it plays a stable role in inhibiting inflammation across all three groups (HC, HPV, and CIN). However, *Lactobacillus iners*, while also showing a negative correlation in these groups, exhibited a more pronounced effect in the HPV and CIN groups compared to the HC group (Figure 4A, C, and E). This suggests that *Lactobacillus iners* may have an increased inhibitory effect on the vaginal microbiome in HPV and CIN patients, specifically by regulating the growth of *non-Lactobacillus* bacterial species.

### Potential Role of *Lactobacillus iners* in HPV Infection

As highlighted by previous research, the percentage of *Lactobacillus iners* tends to increase in patients with HPV infections, but its exact role in the vaginal microenvironment remains unclear. It is difficult to definitively conclude whether *Lactobacillus iners* has a beneficial or detrimental effect in the context of HPV infections. However, our findings suggest that *Lactobacillus iners* may act as a regulatory agent, potentially downregulating the growth of *non-Lactobacillus* species in the vaginal microbiome, thereby exerting a protective or balancing effect in HPV-infected and CIN patients. These results provide important insights into the role of *Lactobacillus iners*, indicating that it may help maintain the stability of the vaginal microbiome by reducing the abundance of more pathogenic, *non-Lactobacillus* bacteria. However, further research is needed to fully understand its dual role and the mechanisms through which it modulates the vaginal environment in the presence of HPV and CIN.

### ELISA Results of Co-cultured Supernatants and Validation with Clinical Samples

To validate the findings from clinical samples, we conducted a co-culture experiment using Hela cells and various bacterial species identified from the heatmap analysis. We collected the supernatants after co-culturing and tested the cytokine concentrations using ELISA. We focused on cytokines that exhibited significant changes in the clinical heatmap results, specifically IL-1α, IL-1β, IL-6, and IL-8. While IL-1α and IL-1β showed a clear negative correlation with *Lactobacillus crispatus* and *Lactobacillus iners* in the clinical samples, the results from the co-culture experiment did not reflect a significant decrease in their concentrations. Moreover, the other bacterial species positively correlated with IL-1α and IL-1β in the clinical samples did not show the expected increase in the lab-based co-culture. This discrepancy suggests that other in vivo factors might be influencing the expression of cytokines and bacteria, which could account for the differences observed between the clinical and lab results (Figure 5A-D). For IL-6 and IL-8, the co-culture results largely mirrored the trend observed in clinical samples, with minimal differences across the bacterial species tested (Figure 5E-H, K, and L). This consistency strengthens the reliability of our findings, especially with regard to these two cytokines, suggesting that the in vitro co-culture system is a valid model for cytokine expression in certain contexts.

In summary, our research examined the correlation between vaginal bacteria and cytokines in Chinese women and revealed several important findings. Firstly, POP patients exhibited higher alpha-diversity, a lower percentage of *Lactobacillus spp.*, and a higher percentage of *non-Lactobacillus* bacteria, suggesting a distinct vaginal microbiome profile compared to healthy controls. Secondly, both *Lactobacillus crispatus* and *Lactobacillus iners* showed a significantly negative correlation with the six cytokines tested, while *non-Lactobacillus* bacteria (e.g., *Gardnerella vaginalis, Sneathia amnii*) exhibited a positive relationship with the same cytokines, indicating their potential role in promoting inflammation. Lastly, *Lactobacillus iners* displayed a significantly negative effect on the vaginal microbiome compared to the healthy control group, suggesting that it may play an inhibitory role in these conditions by reducing the growth of *non-Lactobacillus* species. Our findings contribute to a deeper understanding of the role of vaginal microbiota in gynecological health and inflammation, highlighting potential therapeutic targets for managing conditions like HPV infection and CIN.

## Supporting information

Supplementary statistics

## ACKNOWLEDGMENTS

This work was supported by the NNSF-VR Sino-Swedish Joint Research Programme (82161138017 to W.H.C), Vetenskapsrådet (2021-06112), and Capretto Company (Eurostars 3 project, ID: 2822, project: C125233193), and we would like to thanks for their help and support.

