## Supplementary statistics for "Vaginal Microbiome and Inflammation Among Chinese Women"

### Supplementary Statistis

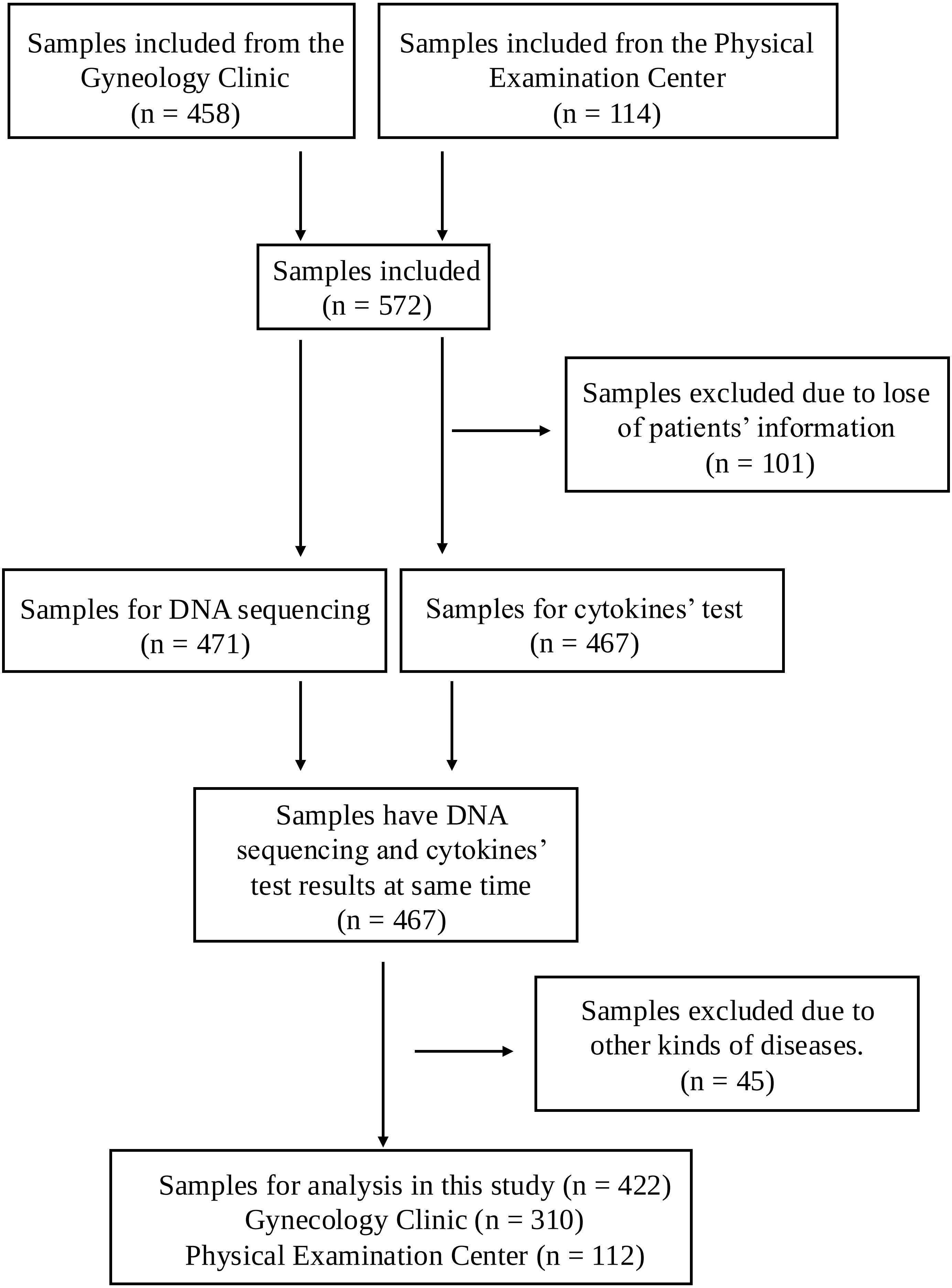


**Overview of clinical samples**

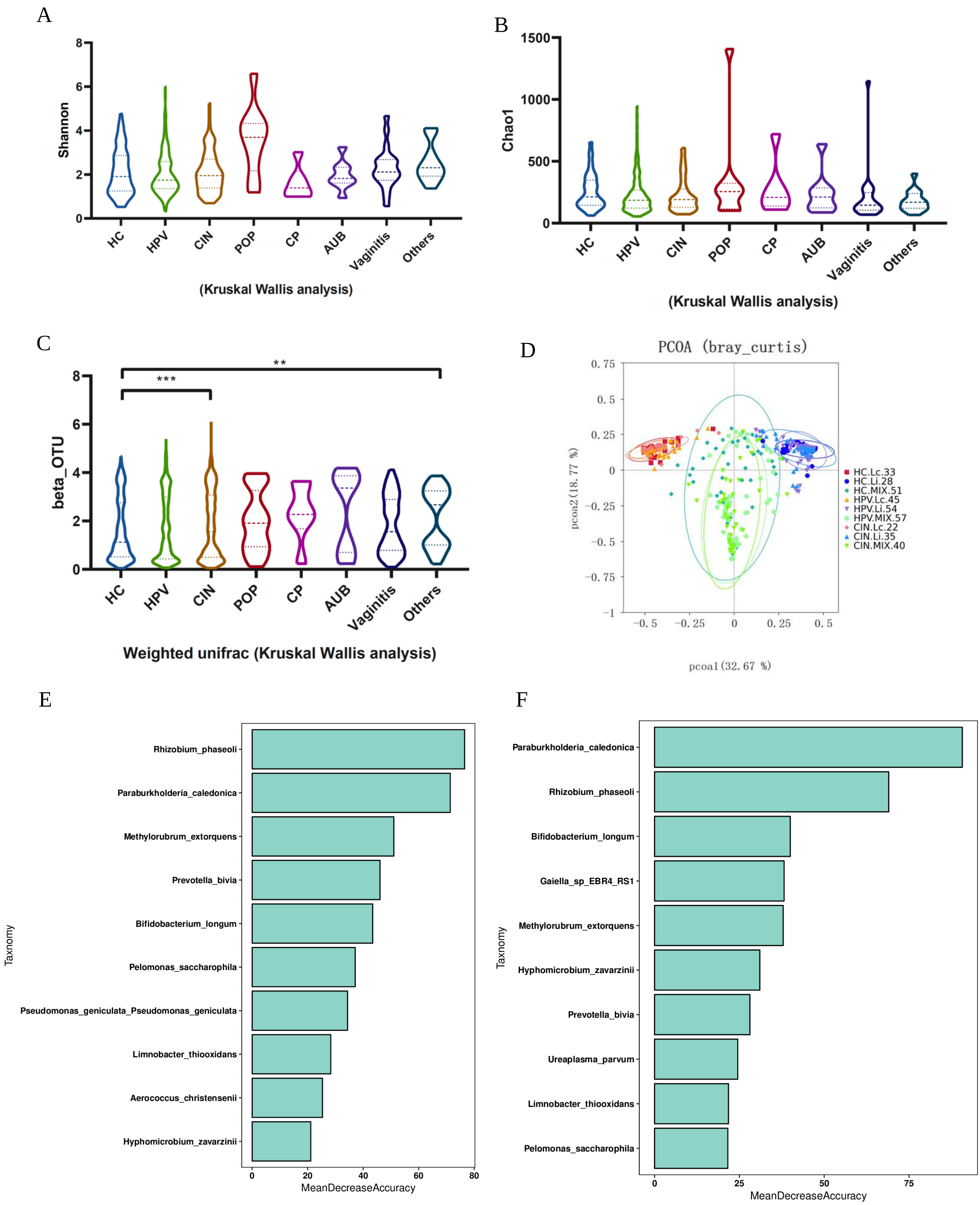


**Supplementray Figure 1: Basic information of 422 clinical samples. A-B)** alpha analysis of 422 clinical samples. **C)** beta analysis of 422 samples. **D)** PCoA result of 422 samples. **E)** Random forest analysis screened top10 species by Mean Decrease of HC and HPV group. **F)** Random forest analysis screened top10 species by Mean Decrease of HC and CIN group. (Kruskal Wallis analysis, * : $p < 0.05$; ** : $p < 0.01$; *** : $p < 0.001$)

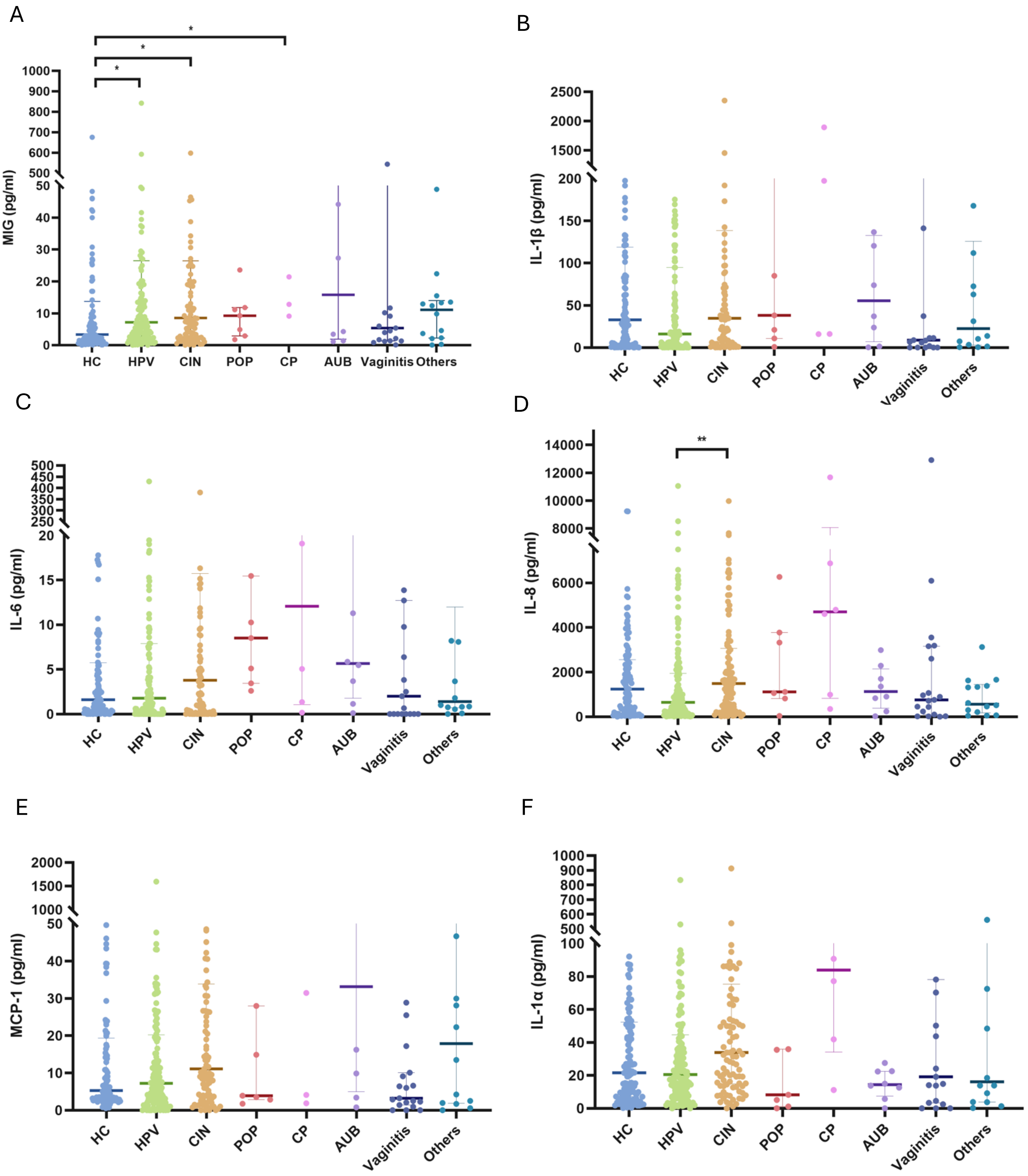


**Supplementray Figure 2: cytokines' expression in HC, HPV, CIN, POP, CP, AUB, Vaginitis, and Others groups. A-F)** Concentrations of MIG, IL-1β, IL-6, IL-8, MCP-1, and IL-1α. (Kruskal Wallis analysis, * : $p < 0.05$; ** : $p < 0.01$; *** : $p < 0.001$)

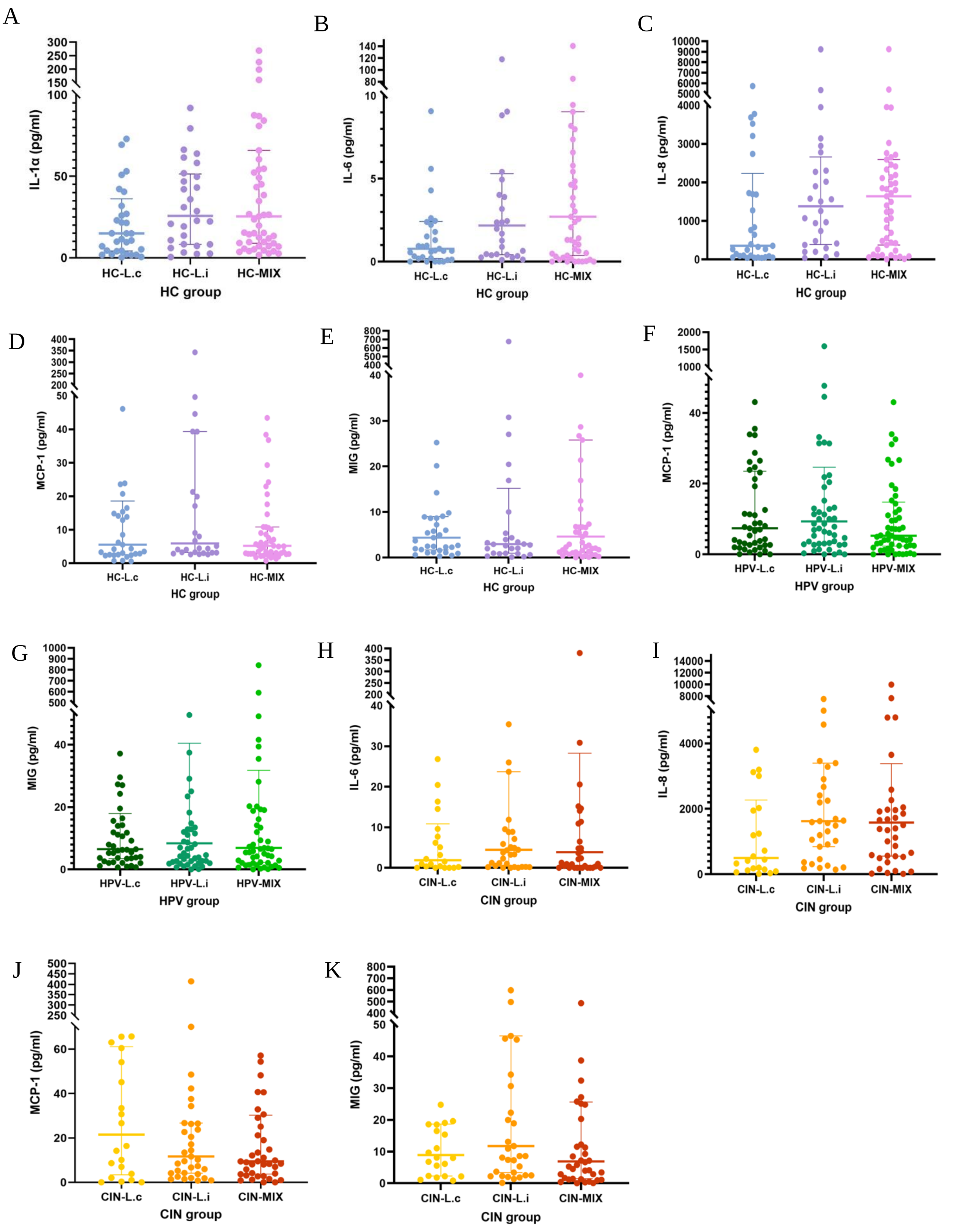


**Supplementary Figure 3: Cytokines' concentration in HC, HPV and CIN groups. A-E)** IL-1a, IL-6, IL-8, MCP-1 and MIG expressed in HC group. **F)** MCP-1 and MIG expressed in HPV group. **H-K)** The concentrations of IL-6, IL-8, MCP-1 and MIG in CIN group. (Kruskal Wallis analysis, * : $p < 0.05$; ** : $p < 0.01$; *** : $p < 0.001$)

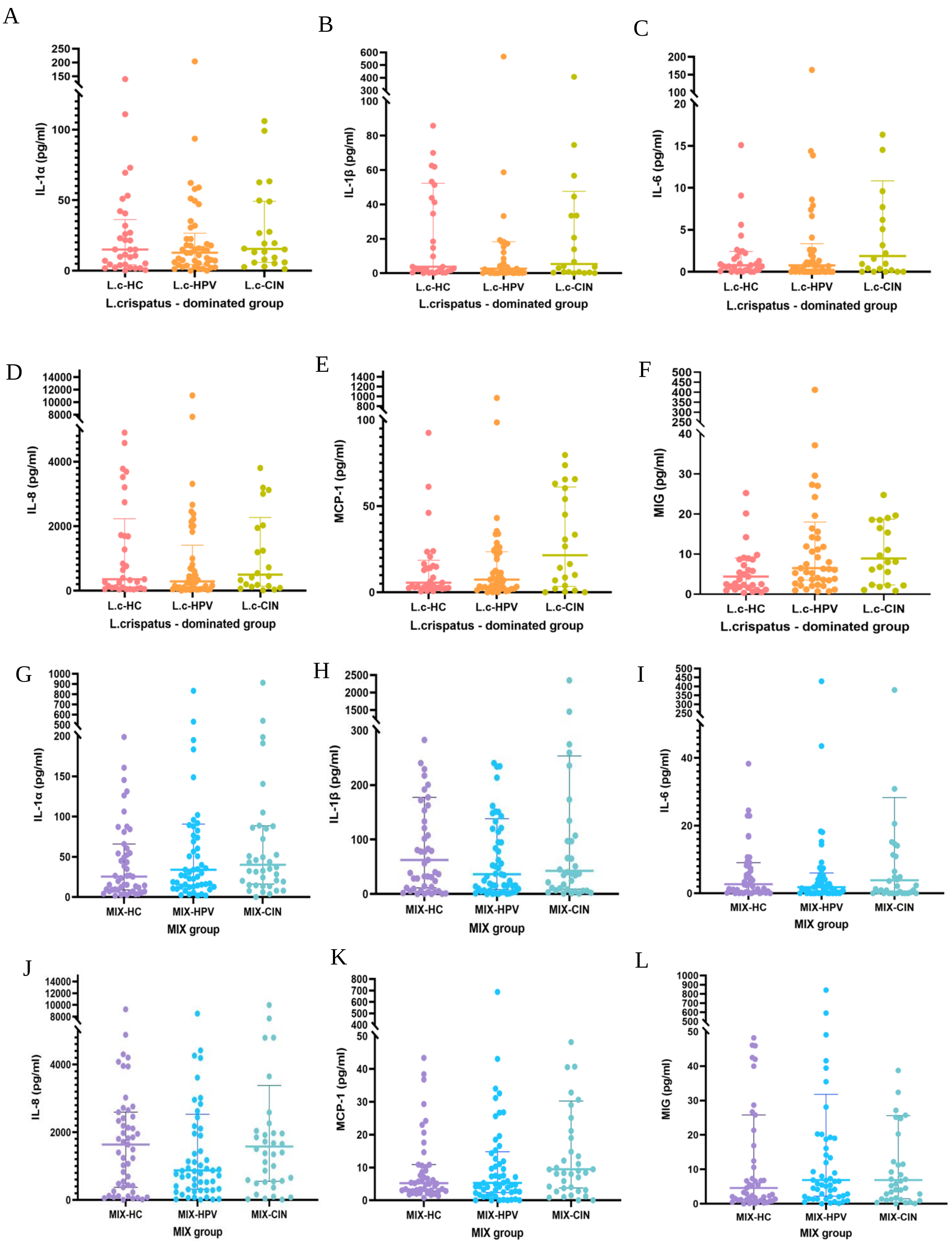


**Supplementary Figure 4: Cytokines' concentration in *L.crispatus*-dominated and MIX groups. A-F)** There were no significant difference of 6 cytokines in *L.crispatus*-dominated group. **G-L)** There were no significant difference of 6 cytokines in Mix group. (Kruskal Wallis analysis, * : $p < 0.05$; ** : $p < 0.01$; *** : $p < 0.001$)

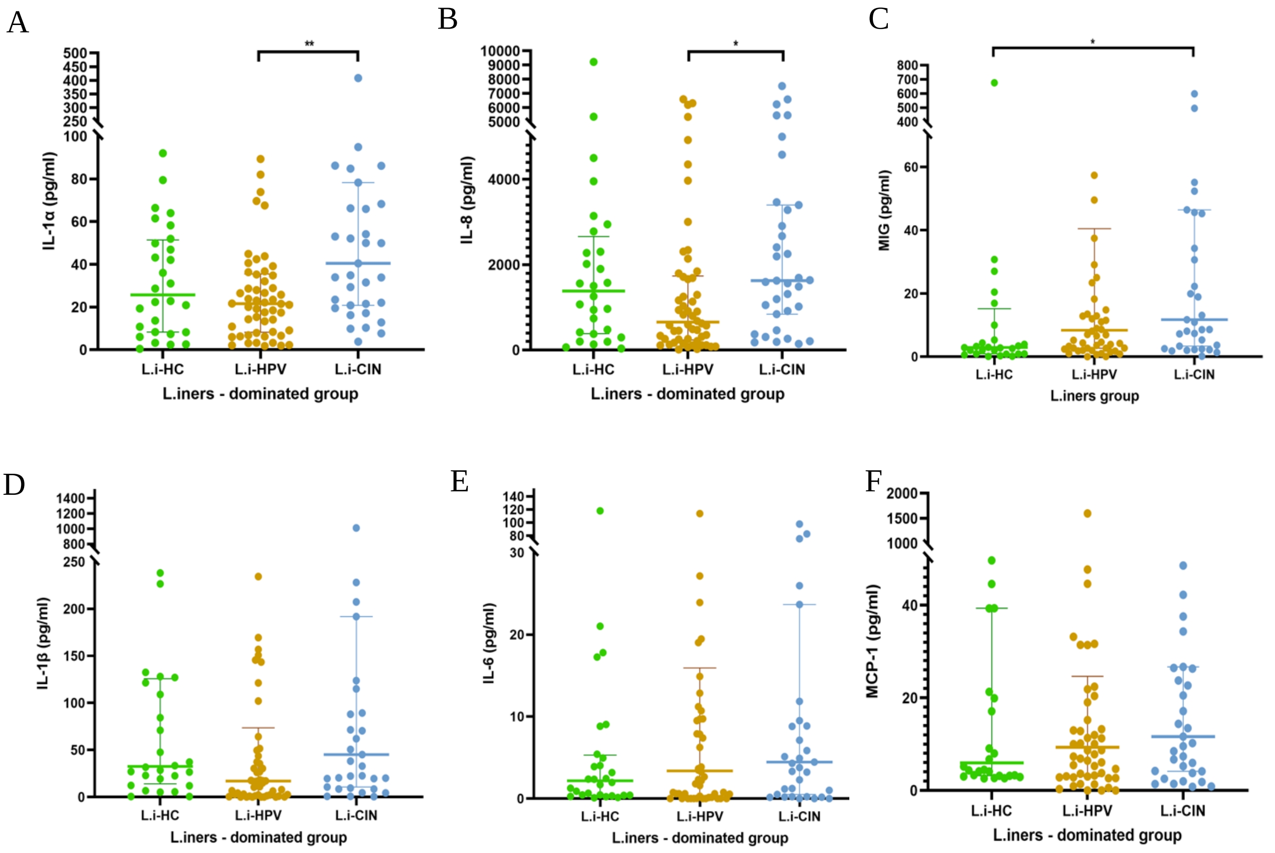


**Supplementary Figure 5: Cytokines' concentration in *L.iners*-dominated group. A-F)** concentratoins of IL-1a, IL-8, MIG, IL-1b, IL-6 and MCP-1 in *L.iners*-dominated group. (Kruskal Wallis analysis, * : $p < 0.05$; ** : $p < 0.01$; *** : $p < 0.001$)
